# Circuit-seq: Circular reconstruction of cut *in vitro* transposed plasmids using Nanopore sequencing

**DOI:** 10.1101/2022.01.25.477550

**Authors:** Francesco E. Emiliani, Ian Hsu, Aaron McKenna

**Affiliations:** Department of Molecular and Systems Biology, Geisel School of Medicine, Dartmouth College, Lebanon, NH 03756, USA; Norris Cotton Cancer Center, Dartmouth-Hitchcock Medical Center, Lebanon, NH 03756, USA

**Keywords:** Long-reads, plasmid, cloning, assembly, sequencing

## Abstract

Recombinant DNA is a fundamental tool in biotechnology and medicine. Validation of the resulting plasmid sequence is a critical and time-consuming step, which has been dominated for the last 35 years by Sanger sequencing. As plasmid sequences grow more complex with new DNA synthesis and cloning techniques, we need new approaches that address the corresponding validation challenges at scale. Here we prototype a high-throughput plasmid sequencing approach using DNA transposition and Oxford Nanopore sequencing. Our method, Circuit-seq, creates robust, full-length, and accurate plasmid assemblies without prior knowledge of the underlying sequence for approximately $1.50 per plasmid. We demonstrate the power of Circuit-seq across a wide range of plasmid sizes and complexities, generating accurate and contiguous plasmid maps. We then leverage our long read-data to characterize epigenetic marks and estimate plasmid contamination levels. Circuit-seq scales to large numbers of samples at a lower cost than commercial Sanger sequencing, accelerating a key step in synthetic biology, with low startup costs make it practical for individual laboratories.

## Introduction

Plasmids are the core building block of recombinant DNA. Researchers can combine and synthesize novel DNA sequences to manipulate cellular biology: from emerging therapeutics like CAR-T cells to synthetic biology circuits that perturb cellular functions (Antebi et al. 2017; Hernandez-Lopez et al. 2021). These constructs are often sensitive to imperfections and require careful sequence validation. Plasmid sequence is typically verified using chain-termination sequencing (also known as Sanger sequencing), first described in 1977 (Sanger, Nicklen, and Coulson 1977). Sanger sequencing requires a complementary oligonucleotide to bind upstream of the sequence of interest, from which random terminated fragments are used to create a consensus sequence (Heather and Chain 2016). However, plasmid sequences are now routinely greater than 10 kilobases, requiring a large number of custom primers, and repetitive regions or imbalanced nucleotide content can be especially challenging for Sanger sequencing (Kieleczawa, Dunn, and Studier 1992). To circumvent these limitations, the field has developed a number of secondary techniques to validate the entirety of the sequence, including restriction enzyme mapping or successive primer walking. The collective reagents and time required to validate individual sequences can halt progress or limit the scope of scientific questions; as we incorporate advances in DNA construction and decreased DNA synthesis costs, we expect these challenges of scale and accuracy to only grow more acute (Gibson et al. 2009; Hughes and Ellington 2017).

Our lab’s need to validate complex plasmid assemblies at scale led us to assess the available methods for high-throughput sequencing and assembly. Recent work has demonstrated the utility of high-throughput plasmid verification using Illumina sequencing technologies (Shapland et al. 2015; Gallegos et al. 2020). The result has been impressive: Gallegos et al. were able to fully assemble a 96-well plate of 2.5 to 3.3 kb plasmids. Unfortunately, these short-read approaches often struggle to achieve high contiguity for more complex plasmids: the same pipeline in Gallegos et al. was only able to assemble 25% of a more complex plasmid pool containing longer repeat elements (Gallegos et al. 2020; Shapland et al. 2015). Sequencing cost is also an issue, as large numbers of plasmids must be pooled to overcome reagent costs, and upfront machine costs are often prohibitive for labs that don’t have direct access to an Illumina machine (Gallegos et al. 2020).

Inspired by a recent publication of a PCR-based plasmid sequencing method (Currin et al. 2019) and the low-cost Nanopore Flongle flowcell, we turned our attention to the Oxford Nanopore long-read sequencing platform. We reasoned that long-read sequences could overcome challenging repeat regions, especially if we were able to preserve the contiguity of the underlying sequence through library generation. This key advantage, coupled with recent advances in sequencing output and computational methods improvement could allow us to accurately assemble even the most complex plasmids. Here we detail our approach, Circuit-seq (**<u>Ci</u>**rcular **<u>r</u>**econstruction of **<u>c</u>**ut **<u>i</u>**n vitro **<u>t</u>**ransposed plasmids), which generates complete maps of synthetic DNA constructs within 24 hours at a cost that is competitive with single chain-terminating sequencing reactions. These assemblies can then be annotated with epigenetic base modifications, and our long-read data can be used to estimate input contamination levels, providing a comprehensive sequence characterization for a wide variety of downstream applications. A Nextflow pipeline is also available implementing the computational aspects of Circuit-seq: https://github.com/mckennalab/Circuitseq (Di Tommaso et al. 2017).

## Results

### Tagmentation leads to robust multiplexing of a diverse set of 96 plasmids

Multiplexing approaches for sequencing have increased throughput and decreased per-sample costs (Shendure et al. 2017). Thus, we set out to develop an approach that multiplexed the capture and assembly of full-length plasmid sequences on a long-read sequencing platform. We initially considered two library generation approaches: (1) restriction digestion of plasmids followed by barcode ligation (**fig. s1**) and (2) barcode tagmentation with the Tn5 transposase (**fig. 1a**) (Reznikoff 2008). While both approaches proved successful, we decided to employ the sequence-agnostic Tn5 transposition technique.

**Figure 1.**
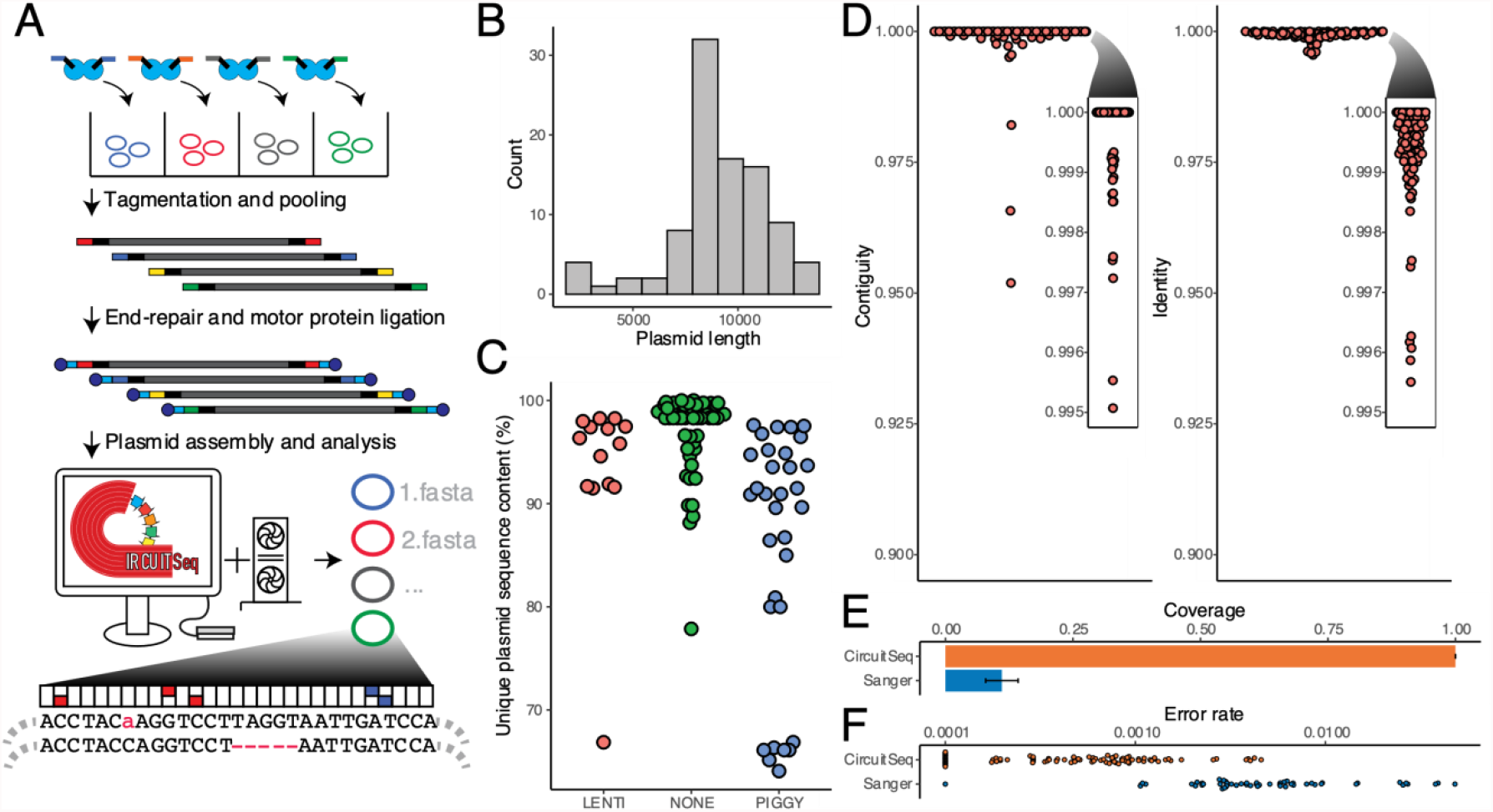
**(A)**. Schematic representation of the Circuit-seq. Plasmids are arranged in a 96 well are tagmented with well-specific barcodes. Samples are then pooled, end-repaired, adapter-ligated, and processed on an Oxford Nanopore Flongle flowcell. The generated data is then run through a custom NextFlow pipeline producing the final assembled sequences. **(B)**. A distribution of the plasmid sizes used in our experiment. **(C)**. Unique (non-repetitive) plasmid rates for our input pool. Lentiviral (LENTI) and piggybac (PIGGY) payload plasmids have more repetitive sequence in their backbones, a key criteria to challenge our assembly pipeline. **(D)**. Contiguity and identity scores for the polished assemblies were calculated by comparison to the known reference. **(E)**. Proportion of the full plasmid covered by Circuit-seq assemblies and Sanger sequencing. **(F)**. The error rate for Circuit-seq assemblies and Sanger sequencing was calculated by comparison to known reference. Sanger sequences were pre-filtered to remove sequences with >10% error to exclude technical errors from the analysis.

The Tn5 enzyme is commonly used for transposase-mediated adaptor insertion and subsequent sequencing (A. Adey et al. 2010). Tn5 is loaded in vitro with an oligonucleotide (oligo) containing the Tn5 mosaic sequence (Reznikoff 2008; A. Adey et al. 2010; A. C. Adey 2021). To uniquely tag individual plasmids, we extended these 19bp mosaic sequences with a set of 96 error-resistant 17bp barcodes from Hawkins et al. (**fig. 1a, table s3**) (Hawkins et al. 2018). Our goal was a limited tagmentation of each plasmid to preserve contiguity and to lower the per-reaction cost. We used limiting quantities of oligonucleotides in an excess of Tn5 to restrict tagmentation, which we tested over a range of concentrations, near the lowest levels used by Picelli et al. (Picelli et al. 2014). Tn5 tagmented libraries were then pooled and prepared using the Oxford Nanopore ligation sequencing kit and loaded onto an individual Flongle flow cell. The results of our optimization (**fig. s2a**) show that our approach is robust over the tested conditions, generating a sizable proportion of full-length reads. Increasing the amount of Tn5 or the duration of tagmentation increased the number of tagmentations per plasmid, though we recovered sufficient fragments from all conditions to generate correct assemblies (**fig. s2b**).

We next tagmented a diverse set of 96 plasmids using the optimized conditions above. Each input plasmid has a known sequence map, ranging in size from 2,433 to 13,286 basepairs (**fig. 1b**). The input plasmids had a wide range of repetitive sequence content (ranging from 0 to 36% of the known map) incorporating both lentiviral and PiggyBac integration vectors (**fig. 1c**). We also used an improved set of Tn5 sample barcodes (v2) with increased hamming distance to improve barcode recovery (**table s4**). We then demultiplexed and processed individual plasmids with a custom NextFlow pipeline (Di Tommaso et al. 2017). Reads assigned to plasmid barcode IDs accounted for 81% of reads, resulting in 150-4619x (median 1118) coverage of our plasmid set, with little correlation to input plasmid length (r = −0.16, **fig. s3a**). As expected, we observed peaks in fragment sizes at known plasmid lengths, with increasing proportions of smaller fragments which are presumably from DNA fragmentation in library preparation and sequencer length bias (**fig. s3b**) (Amarasinghe et al. 2020). Consistent with this hypothesis, we observe more full-length fragments in smaller plasmids (**fig. s3c**). We then filtered and corrected each sample’s reads to generate error-corrected consensus sequences where the distribution more closely matches single and double tagmentation events (**fig. s3d**) (Koren et al. 2017).

### De novo plasmid assembly and validation

We then assembled plasmids de novo from the corrected reads, avoiding any biases from reference-based assembly approaches (Lunter and Goodson 2011). We used a two-pronged assembly approach, leveraging both Flye and Miniasm long-read assemblers to generate competing initial consensuses from which the longest single-contig assembly is taken forward (R. R. Wick and Holt 2019; Kolmogorov et al., n.d.) (https://github.com/lh3/miniasm). This is followed by subsequent rounds of refinement with Medaka and custom post-processing steps to eliminate whole genome duplications and other common assembly artifacts (https://github.com/nanoporetech/medaka). Our pipeline was able to assemble 95 of 96 plasmids to a 100.00% median contiguity and 99.95% identity (**fig. 1d**), an improvement over existing Illumina sequencing approaches (Gallegos et al. 2020). No reads for the missing assembly were recovered from the raw data, suggesting that it was due to technical error during sample preparation.

We next wanted to determine the accuracy of our approach compared to conventional validation techniques. We collected 64 Sanger sequencing reactions from 36 different plasmids within our Nanopore assembly dataset. We converted the resulting trace files and filtered them by quality score using Mott’s algorithm to clip low-quality regions (Ewing et al. 1998). We further removed seven Sanger sequencing results where the error rate was above 10% to avoid biasing our results from potential user error (**fig. s4e and f, methods**). 57 remaining Sanger sequencing results covered a median of 10.3% of their respective plasmid with an average aligned length of 1023 bases (**fig. 1e**). The Sanger median per-base error rate was 0.4%, approximately a 10-fold higher rate than our Nanopore assembly approach (**fig. 1f**). In aggregate, our pipeline successfully assembled high contiguity and identity maps even when faced with highly repetitive sequences.

### Plasmid epigenetics

Given the success of our assembly pipeline, we wondered if we could use our rich sequencing data to provide validation that is inaccessible from chain-terminating sequencing results. Bacterial epigenetic patterns are carried over to plasmid sequences; both *dcm* and *dam* methylation can be transferred to matching recognition sites in plasmids propagated by *E. coli* (Marinus and Løbner-Olesen 2014). This methylation has a number of implications for synthetic biology, including methylation-sensitive restriction enzymes, the stability of repeat sequences, and downstream sensitivity issues in the target organism (Kolek et al. 2016; Nichol and Pearson 2002). Given Oxford Nanopore’s ability to directly capture DNA modifications, we were interested to see if we could profile the epigenetic status of our assemblies. We then extended our pipeline to call methylation modifications present in each plasmid assembly. As initial validation of this approach, we created consensus DNA sequence logos from the 6 methyladenine (6mA) and 5 methylcytosine (5mC) patterns detected in our 96-plasmid run, recovering known sequences of *dam* (GATC) and *dcm* (CCYGG) (**fig. 2a**).

**Figure 2.**
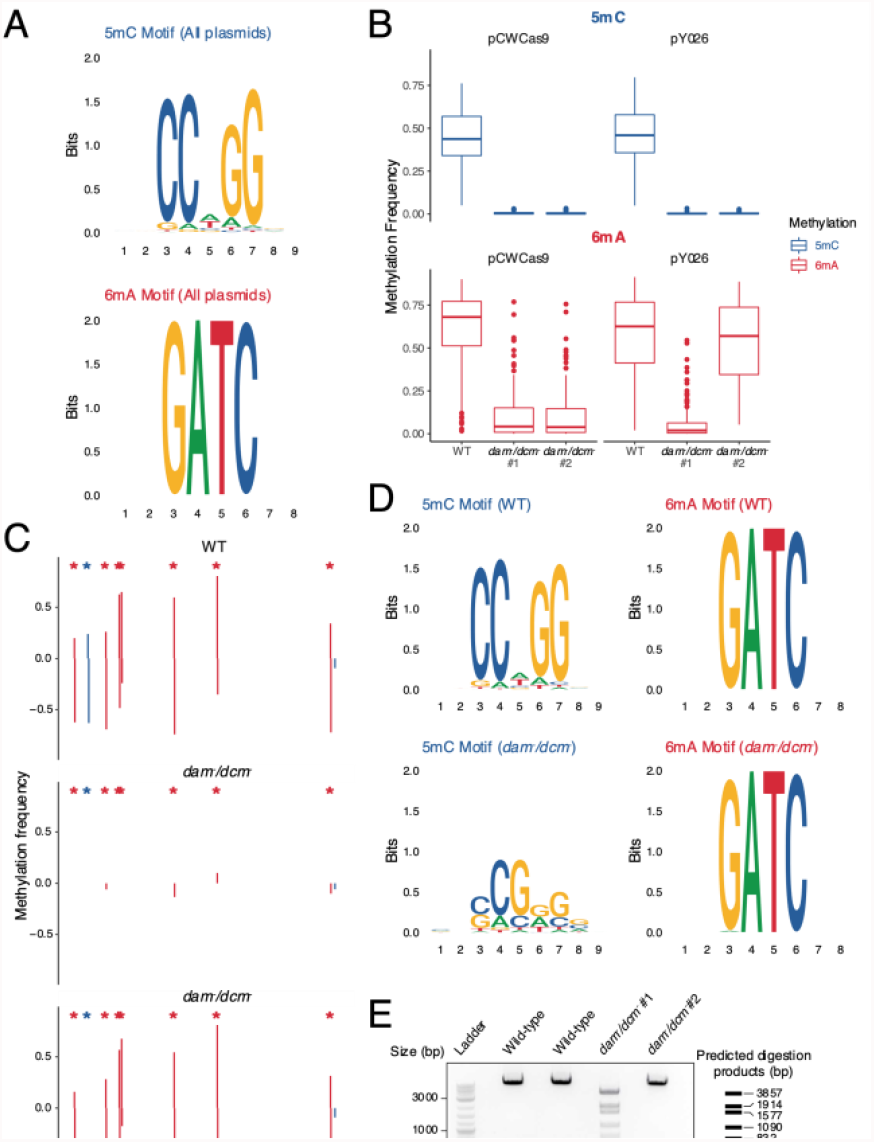
**(A)** DNA sequence logos generated from plasmid methylated 5-methylcytosine (5mC) and 6-methyladenine (6mA) call sequences representing known dcm and dam consensus sequences, respectively. **(B)** Global 5mC and 6mA rates from plasmids grown in conventional *dcm*^*+*^*/dam*^*+*^ *E. coli* (WT) and *dcm*^*-*^*/dam*^*-*^ *E. coli* (**methods**). **(C)** Visualizing methylation frequencies across a region of pY026 plasmid shows methylation localizing to consensus motifs (asterisks) in pairs, corresponding to the top and bottom strands. And highlights the complete loss of 5mC methylation in the *dcm*^*-*^*/dam*^*-*^ samples, and the partial and complete conversion to *dam*^*+*^ in the first and second *dcm*^*-*^*/dam*^*-*^ samples, respectively. **(D)** DNA sequence logo plots from methylation sites of plasmids grown in WT and *dcm*^*-*^*/dam*^*-*^ *E. coli* showing the loss of 5mC but recovery of 6mA. **(E)** Validation of methylation using 6mA sensitive BclI digestion of MluI linearized pY026 plasmid. BclI digestion failed to cleave the methylated plasmid as well as the *dam*^*+*^ recovered plasmid, but near-complete digestion of the *dam*^*-*^ plasmid. Predicted digestion products annotated on the right.

To further validate the accuracy of our approach we then transformed plasmids into both NEB Stable Competent cells (WT) and *dam*^*-*^*/dcm*^*-*^ competent cells that lack both methyltransferases. One WT plasmid and two separate colonies of *dam*^*-*^*/dcm*^*-*^ plasmids were processed according to our normal Circuit-seq protocol. In the *dam*^*-*^ */dcm*^*-*^ plasmids, methylation calls at motif sequences show a drastic drop of 6mA and a near-complete loss of 5mC (**fig. 2b**). The low, but not abolished, levels of 6mA can be explained by spontaneous re-activation of *dam* through the loss of the knockout transposon sequence (New England Biolabs n.d.). This is evident in the second replicate of pY026 *dam*^*-*^*/dcm*^*-*^ which has a near-complete reactivation of *dam* but a complete absence of *dcm* activity. Highlighting a stretch of this pY026 assembly, we see a mirroring of methylation patterns on each of the positive and negative strands, confirming that the regained *dam* activity is consistent with true methylation patterns (**fig. 2c**). As can also be seen in the DNA sequence motifs from the various conditions, there was no clear sequence motif enrichment in the 5mC motif in the *dam*^*-*^*/dcm*^*-*^ samples, while the 6mA successfully recreates the dam motif (**fig. 2d**). Finally, to experimentally validate the methylation levels, we digested plasmids with a *dam*-sensitive restriction enzyme, BclI. The resulting gel shows near-complete digestion of the first *dam*^*-*^*/dcm*^*-*^ plasmid but no visible fragments in the second colony, validating the computational characterization (**fig. 2e**).

### Contamination detection

Plasmid preparations can be contaminated by bacterial DNA or other passenger plasmids that confound downstream experiments. Given the number of sequencing reads produced for each plasmid, we wondered if we could leverage this read depth to estimate potential contamination levels. Given the shared components of many plasmids, i.e. the origin of replication or lentivirus components, we reasoned that previously developed SNV-based contamination estimation methods would be ill-suited for the task (Cibulskis et al. 2011; Jun et al. 2012). Instead, we used a simple Bayesian model to leverage the proportion of unaligned bases (‘clipped’ bases) as well as unmapped reads during sequence alignment, taking into account the shared nature of plasmid components (**fig. 3a, methods**).

**Figure 3.**
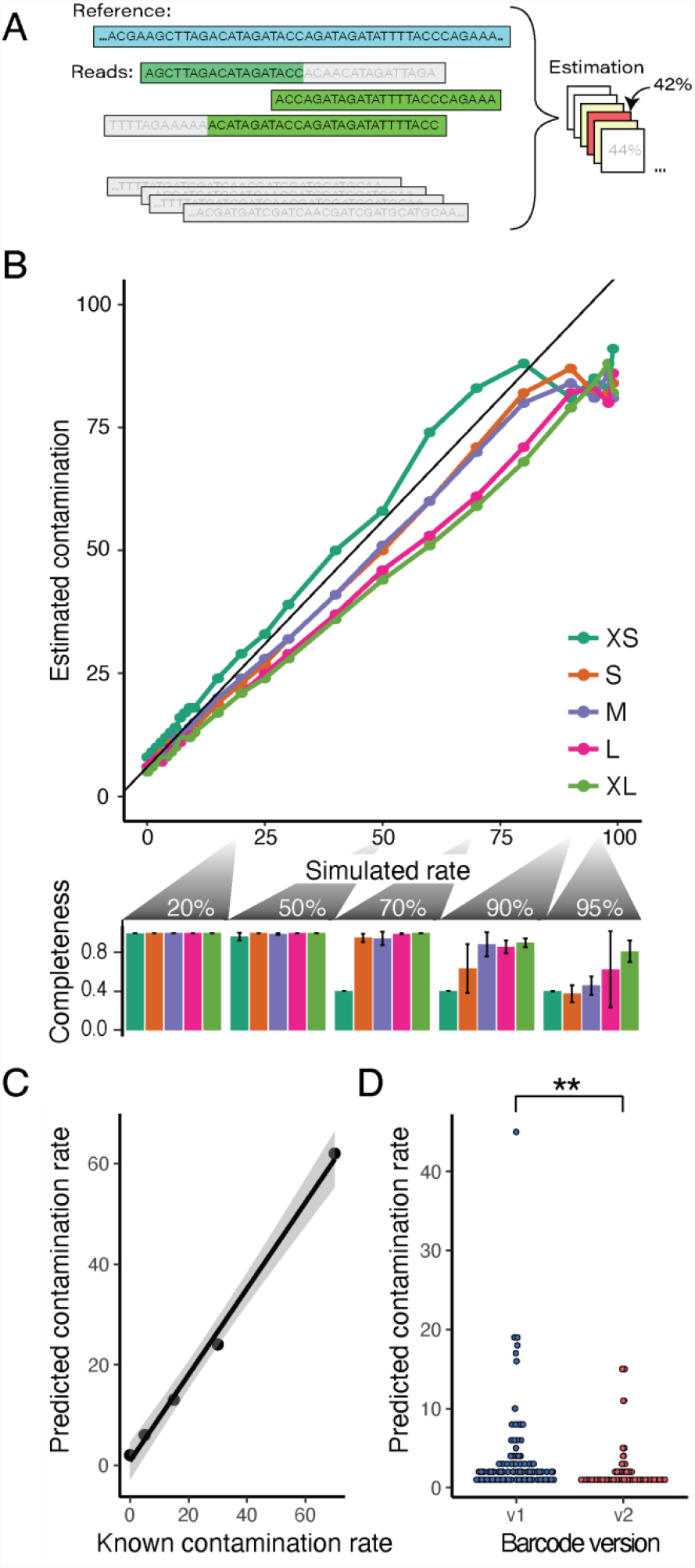
**(A)** Contamination rates are calculated from the proportion of unaligned bases and the global shared sequence proportions between plasmids, estimated over a range of contamination hypotheses (**methods**). **(B)** In silico contamination estimates of assembled Addgene plasmids using simulated reads over a range of plasmid lengths (upper panel, XS=extra small to XL=extra large maps). Our estimates track the programmed rates well (r = 0.97, RSME = 9.1), though at higher contamination levels our assemblies had issues that result in poor contamination estimates, seen in the assembly completeness drop off for small plasmids at >70% contamination and at >90% for larger plasmids (lower panel). **(C)** Correlation of an in vitro mixing experiment of two known plasmids prior to sequencing and assembly. Contamination estimates were well correlated to their mixing proportion (r = 0.997, RSME = 4.66) though we failed to assemble the plasmids at the two highest contamination levels (85 and 95% contaminated). **(D)** Estimated contamination rates significantly dropped with our improved Tn5 barcoding design (p = 0.00119).

We first validated our approach using computationally generated reads from commonly used plasmids available from the Addgene repository. Five plasmids were chosen spanning a range of sizes and were subsequently contaminated by random reads from 91 other constructs at set levels. The simulated reads were then used as input to our existing pipeline and our computational tool was used to predict contamination levels (**fig. 3b, top**). Contamination levels were relatively well-predicted by our simple method (r = 0.97, RSME = 9.1), though were less accurate at high contamination levels due to underlying assembly failures (**fig. 3b, bottom, fig. s4a**). When we estimated contamination using known plasmid maps instead of our assemblies, we saw similar results, with more consistent estimates at higher contamination levels (r = 0.99, RSME = 7.2, **fig. s4b**).

To further validate our contamination pipeline, we experimentally mixed one plasmid preparation into another over a range of contamination rates. We then sequenced these mixtures in individual wells in an additional Circuit-seq run. Our two most contaminated plasmids, 85% and 95% contaminated, failed to assemble, but computational estimates in the remaining samples tracked the known experimental rates (r = 0.997, RSME = 4.66) (**fig. 3c**). We then profiled overall contamination rates in both our v1 and v2 barcoding runs (**fig. 3d**). Contamination levels were correlated with assembly errors (Pearson’s r = 0.32), suggesting an avenue to further improve assembly results.

## Discussion

Here we detail our new high-throughput technique, Circuit-seq, an end-to-end plasmid validation pipeline that leads to near-perfect assemblies. This technique, complementing other 2nd and 3rd generation sequencing approaches, provides a comprehensive map of both simple and complex plasmids. In comparison to existing Illumina-based techniques, our long-read approach can assemble through large repetitive regions, as well as to characterize epigenetic marks leveraging the unique advantages of the Nanopore platform. We are then able to couple this with computational tools to provide end-users a comprehensive view of the resulting plasmid with errors rates an order of magnitude lower than Sanger sequencing.

Our assemblies are also cost-effective. The reagents required per 96-well run are approximately $140 (or < $1.50 per plasmid), significantly cheaper than commercial Sanger sequencing (**table s1**). Additionally, the low start-up cost of the Oxford Nanopore platform allows researchers with even moderate validation needs to consider this approach. Higher density barcoding (for instance 384-well plates in parallel) would further reduce costs. Additionally, Circuit-seq can be run with the larger Minion flow cells that produce data at 5-10x the speed, reducing the amount of time required to obtain assemblies. The Circuit-seq protocol is relatively straightforward once the reagents are obtained, and new users in our lab have successfully generated data in their initial experiment. To further enable the adoption of this technique, we’ve made all barcode sequences and computational tools publicly available.

In the future, we hope to adopt new computational and experimental improvements on the Oxford Nanopore platform into Circuit-seq, including higher accuracy Nanopore kits and flowcells. It would be straightforward to include these improvements as they become available for the lower-output Flongle, further decreasing error rates and potentially solving our limited remaining contiguity challenges. We have also had preliminary success directly sequencing plasmids from bacterial colonies, replacing traditional colony PCR, though optimization is still needed to increase consistency and yield.

We have found Circuit-seq to be of immense practical value. This is especially true for projects with highly repetitive or complex sequences, where Sanger sequencing results were hard to interpret or outright failed. Even in routine experiments, the effort of Circuit-seq is rewarded when we discover a plasmid backbone mutation or mislabeled tube that would have gone undiagnosed until much later in the project. Combined with the competitive cost, we are excited by the prospect of Circuit-seq or other modern techniques supplanting Sanger sequencing, providing a single-pass, comprehensive validation of DNA constructs.

## Acknowledgements

We would like to thank the members of the McKenna lab for experimental help, advice, and for providing the stream of complex plasmids needed to validate this approach. An additional thanks to the members of the Dartmouth community who also contributed plasmids in the early development phase. We would especially like to thank Rachel Saxe and Maryam Fathi for testing both the protocol and computational pipelines. This work was supported by funding from the Neukom Insitute at Dartmouth College and The Norris Cotton Cancer Center (NCI 5P30CA023108-37). A.M. is supported by NIH/NHGRI (R00 HG010152-04), the Pew Biomedical Scholars Fellowship, and the V Foundation. F.E. was supported from the Tenney Fellowship and I.H. from the Neukom Insitute and the Sophomore Research Scholarship at Dartmouth College.

## Methods

### Restriction digest approach

We created six test ligation adapters with unique 20 basepair barcode identifiers (**table s2**), containing XhoI, EcoRI, XbaI, BmtI, BclI, and SacI restriction sites. Restriction sites were chosen by scanning a listing of all Addgene plasmid sequences, removing sequences shorter than 2000 bases, and generating a cover-set of enzymes that would maximize the number of sequences with at least one single or double-cut enzyme location (**fig. s1**). A common forward and a unique reverse adapters were annealed at equal molar ratio and extended with a single cycle of Kapa HiFi polyermase using the manufacturer recommended conditions and an extension time of 5 minutes. Adapters were then cut with the target restruction enzyme and purified with Zymo Research Clean & Concentrate kit. Target plasmids were processed similarly in parallel. Adapters were then ligated to their corresponding plasmid at 3x molar excess using New England Biolab’s Quick Ligation kit for 10 minutes at room temperature, followed by a 0.5X Ampure cleanup to remove excess Adapters, followed by standard Oxford Nanopore ONT LSK-109 or ONT LSK-110 ligation protocols and loaded onto a Oxford Nanopore Flongle flowcell.

### Tn5 barcode design

Our 17-nucleotide (nt) barcodes were obtained from the Hawkins et al. using a set with a minimum of 2-error corrections (Hawkins et al. 2018**)(table s3)**. For our version 2 barcodes (v2) we used 27nt barcode sequences, combing Finkelstein’s 13nt and 14nt with 2 error corrections were randomly sampled from the Finkelstein et al. sets, combined, and analyzed using the R package DNABarcodes. We resampled 10,000 combinations and selected an optimized set with a median hamming distance of 30 and a guaranteed error correction of 7bp (Buschmann and Bystrykh 2013). The resulting primers were then screened with Integrated DNA Technologies (IDT) OligoAnalyzer to select 96 primers with optimal delta G > −4kcal/mole (**table s4**). Barcodes were annealed with a common phosphorylated oligo (/5Phos/CTGTCTCTTATACACATCT) by heating to 95°C for 5 minutes and slowly cooling to room temperature.

#### Tn5 purification and storage

Tn5 was purified by the University of California Berkeley Quantitative Bioscience MacroLab Core Facility as per previously described protocols (Hennig et al. 2018). A long-term storage stock solution of Tn5 at 4mg/mL was kept at −80°C in storage buffer (50mM Tris-HCl pH 7.5, 800 mM NaCl, 0.2 mM EDTA, 2 mM DTT, 10% glycerol). A stock solution at 40ug/mL Tn5 was kept at −20°C in the working buffer (50mM Tris-HCl pH 7.5, 800 mM NaCl, 0.2 mM EDTA, 2 mM DTT, 50% glycerol).

### Tagmentation reactions

For 96 plasmid on an r9.4 Flongle: 50ng of plasmid was combined with 2.5pmoles of annealed barcode oligo and 40ng of Tn5 in reaction buffer (50mM Tris-acetate pH 7.5, 150mM potassium acetate, 10 mM magnesium acetate, 4 mM spermidine, 1mM DTT) in a 5ul reaction. For runs with fewer than 48 plasmids, all quantities can be doubled. Using a thermocycler, samples were incubated at 23°C for 10 minutes, 37°C for 10 minutes, and then 55°C for 5 minutes. The reaction was stopped with 1 volume of 0.2% SDS for 5 minutes at 23°C. Samples were pooled and cleaned up with 0.5x volume of Ampure beads according to established protocols and eluted in 25ul of water. For the r10.3 minion run all volumes were doubled but the rest of the protocol remained the same.

### Oxford Nanopore (ONT) library preparation

Plasmids sequences are available for our V1 and V2 efforts in **supplemental tables 5 and 6**. Samples were prepared in accordance with the ONT LSK-110 manual for Flongle libraries. In brief, the purified tagmented libraries were repaired using a combined NEBNext FFPE DNA Repair Mix and NEBNext Ultra II End Repair/dA-Tailing Module by incubating at 20 °C for 7.5 minutes and 65°C for 7.5 minutes. Samples were purified with 0.5x vol of Ampure beads and eluted into 30ul of water. Ligation was performed with NEB Quick Ligase, ONT ligation buffer, and Oxford Nanopore’s adapter mix (AMX) for 10 minutes at room temperature. Samples were purified with 0.5x Ampure beads and then washed with ONT long fragment wash buffer instead of 70% ethanol, and eluted into 7ul of ONT elution buffer. 5ul of the resulting library was loaded into a Flongle as per ONT specifications.

### Demethylated plasmid preparation

Plasmids were transformed into NEB *dam-/dcm-* competent *E. coli* (C2925I) as per the vendor’s protocols and plated onto LB-agar with ampicillin. The following day we picked two colonies per plasmid and grew overnight in LB-Broth with ampicillin for plasmid extraction with the Qiagen Miniprep kit.

### Data Generation and Assessment Workflow

We established a NextFlow pipeline using a prepackaged Docker container to facilitate adoption and reproducibility (Di Tommaso et al. 2017). This pipeline performs all computational steps from base calling to assembly polishing with the ability to create assembly statistics if reference sequences are provided. The pipeline and related documentation are available on our Github (https://github.com/mckennalab/Circuitseq).

Basecalling was performed with ONT guppy software version 5.0.16+b9fcd7b5b using the r941_min_sup_g507 mode. When necessary the pipeline parameter file can be modified to work with different basecalling models or flow cells. Fastq files are then binned using Oxford Nanopore’s Guppy demultiplexing function using a custom barcode configuration file. Porechop (Pryszcz and Novoa 2021; R. Wick et al. 2017)(https://github.com/rrwick/Porechop) is then used to trim adaptor sequences and discard chimeric reads resulting from aberrant ligation products. Sequences shorter than 500bp are filtered with nanofilt (De Coster et al. 2018)(https://github.com/wdecoster/nanofilt). Filtered reads are corrected with Canu (Koren et al. 2017)(https://github.com/marbl/canu) and assembled with miniasm (https://github.com/lh3/miniasm) and Flye assemblers (Kolmogorov et al., n.d.). The assembly is then polished with one round of Medaka (https://github.com/nanoporetech/medaka) to reduce error, which increases the fidelity of duplication removal using DupScoop (https://github.com/mckennalab/DupScoop), a tool we developed to resolve a common error in Flye assemblies where the assembly is perfectly duplicated. The assemblies are then rotated 50% of their length. We found that the edges of the circular assemblies do not get polished as effectively, and rotating the assemblies allows these sequences that are now in the center of the assembly to be polished efficiently during the following three rounds of Medaka. If a reference is provided, the assemblies undergo assessment of identity and contiguity (R. R. Wick and Holt 2019), which generate the values used within the paper. We then define each assembly’s ‘completeness’, ranging from [0,1], as the value:

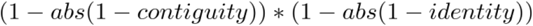

This is a somewhat conservative calculation, given that contiguity and identity values are not fully independent.

### Methylation calling and analyisis

The raw fast5 reads are demultiplexed using the Guppy demultiplexed fastq files using fast5_subset from ont_fast5_api (https://github.com/nanoporetech/ont_fast5_api). The fast5s are then basecalled using Guppy version 4.4.2+9623c16 with the methylation-trained model dna_r9.4.1_450bps_modbases_dam-dcm-cpg_hac.cfg. The methylated sites are then detected and annotated using modPhred (Pryszcz and Novoa 2021)(https://github.com/novoalab/modPhred). For the purpose of creating unbiased DNA sequence logos modPhred was run to detect minimum modification frequencies of 0%, to include all sites that were modified across all replicates. To compare methylation at frequencies between plasmids grown in WT vs *dam*^*-*^ */dcm*^*-*^ competent *E. coli*, we restricted analysis to sites with consensus methylation sequences.

### Contamination - computational simulation of known plasmids

We simulated reads from established plasmids maps downloaded from Addgene. Our goal was to sample from commonly used plasmids, so we first took sequences from the Addgene ‘top 15’ list, with the remaining 84 randomly selected by traversing the Addgene ‘Blue Flame Award’ list alphabetically and selecting a single Blue Flame plasmid from each lab (https://blog.addgene.org/15-years-of-addgene-the-top-15-plasmids) attempting to avoid any related plasmids. The full list of the simulated plasmids is in **supplemental table 7**. The resulting plasmid lengths ranged from 2830 to 16973 bases, with a mean of 8080. Plasmid sequences were downloaded and reads simulated using a version of BadRead customized to empirically draw read lengths from our established fragment lengths over a circular reference, using standard BadRead parameters for ‘bad’ reads (junk reads and chimeric reads) totaling 3% (R. Wick 2019) (https://github.com/aaronmck/Badread). Contamination was simulated for the five Addgene plasmids chosen as our controls (Addgene IDs 31815, 16337, 12251, 49792, and 52961) in five replicates each, representing relatively small to relatively large plasmids. These control plasmids were artificially contaminated with reads randomly drawn from all other plasmids at the following contamination rates: 0, 1, 2, 3, 4, 5, 6, 7, 8, 9, 10, 15, 20, 25, 30, 40, 50, 60, 70, 80, 90, 95, 98, and 99%.

Contamination is then estimated by first aligning Nanopore reads to a duplicated reference map of the plasmid (to capture reads spanning the circular breakpoint) using Minimap2 (https://github.com/lh3/minimap2). Both aligned and unaligned reads are then assessed using a custom Python script. Our Bayesian formulation assumes a flat prior across equality spaced, discrete contamination level hypotheses, much like our approach in Cibulskis and McKenna et al. (Cibulskis et al. 2011):

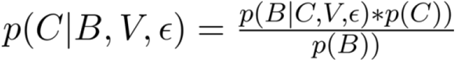

Where *p*(*C*)is our flat prior and the denominator *p*(*B*) is the same over all contamination (*C*) levels. We then need to evaluate the likelihood function. Assuming reads are independent and that our randomly-sampled bases from the read-alignment pair are independent, controling for read length, we can define the likelihood as a product of probabilities:

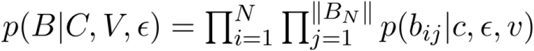

With i..N representing the individual sequencing reads. We can define the following piecewise function:

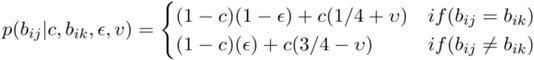

Where b_ij_ and b_ik_ represent the corresponding aligned reference base j and sequencing base k when comparing read i; c is the contamination rate, and epsilon is the Phred-scaled error rate. The parameter v represents a constant shared sequence proportion between plasmids, estimated by comparing kmer proportions between all sampled addgene plasmid sequences (here set to 0.1984). Code to estimate this parameter is included in the GitHub repository. Contamination likelihoods are then normalized to 1 over the range [0,1] and the mode of this distribution is found representing the maximum a posteriori (MAP) score. For simplicity in this paper we chose to sample over 100 bins from [0,1], though increasing this resolution is trivial at the cost of computational time.

## Supplementary Figures

**Sup. figure 1.**
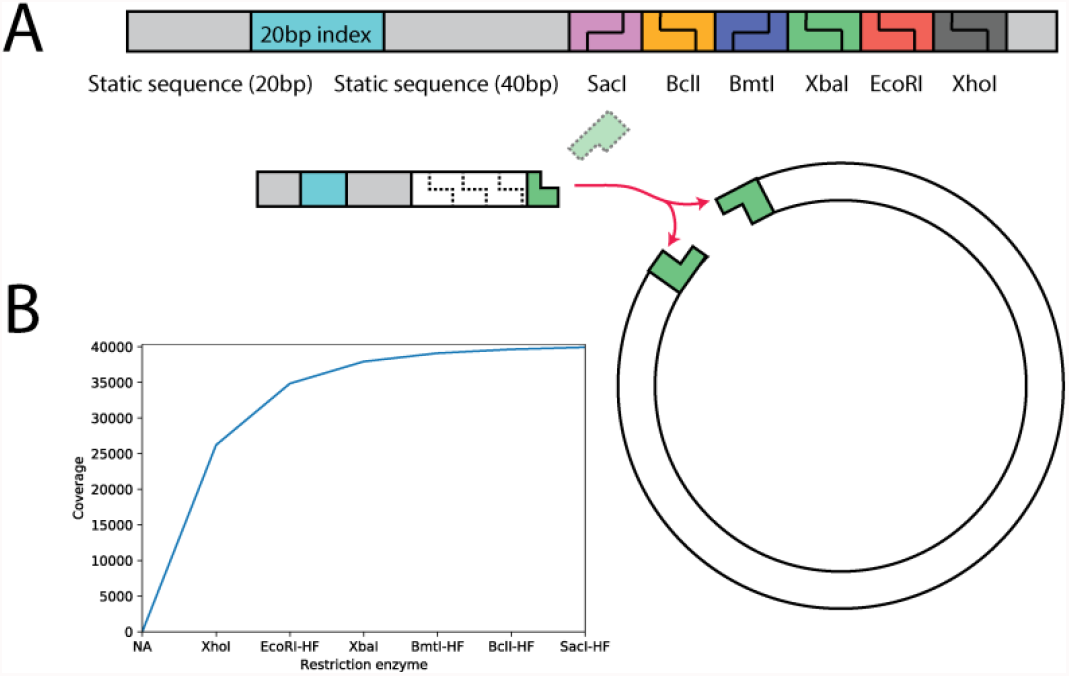
**(A)**. Barcode ligation requires a single, known restriction site (**green**) for each plasmid that is shared with the ligation adapters. Using a set cover approach, we searched all known plasmids in the Addgene database to establish a candidate of restriction enzymes that cut once and only once per plasmid (**see methods**). From this search, we chose a set of 6 restriction enzymes that were incorporated into unique tagged barcodes for ligation. **(B)** These restriction enzymes covered a total of 62% of the 63,017 Addgene plasmids with a single restriction site. Although this approach offers a successful and streamlined approach to barcoding our hope was to be agnostic to the sequence of each plasmid.

**Sup. figure 2.**
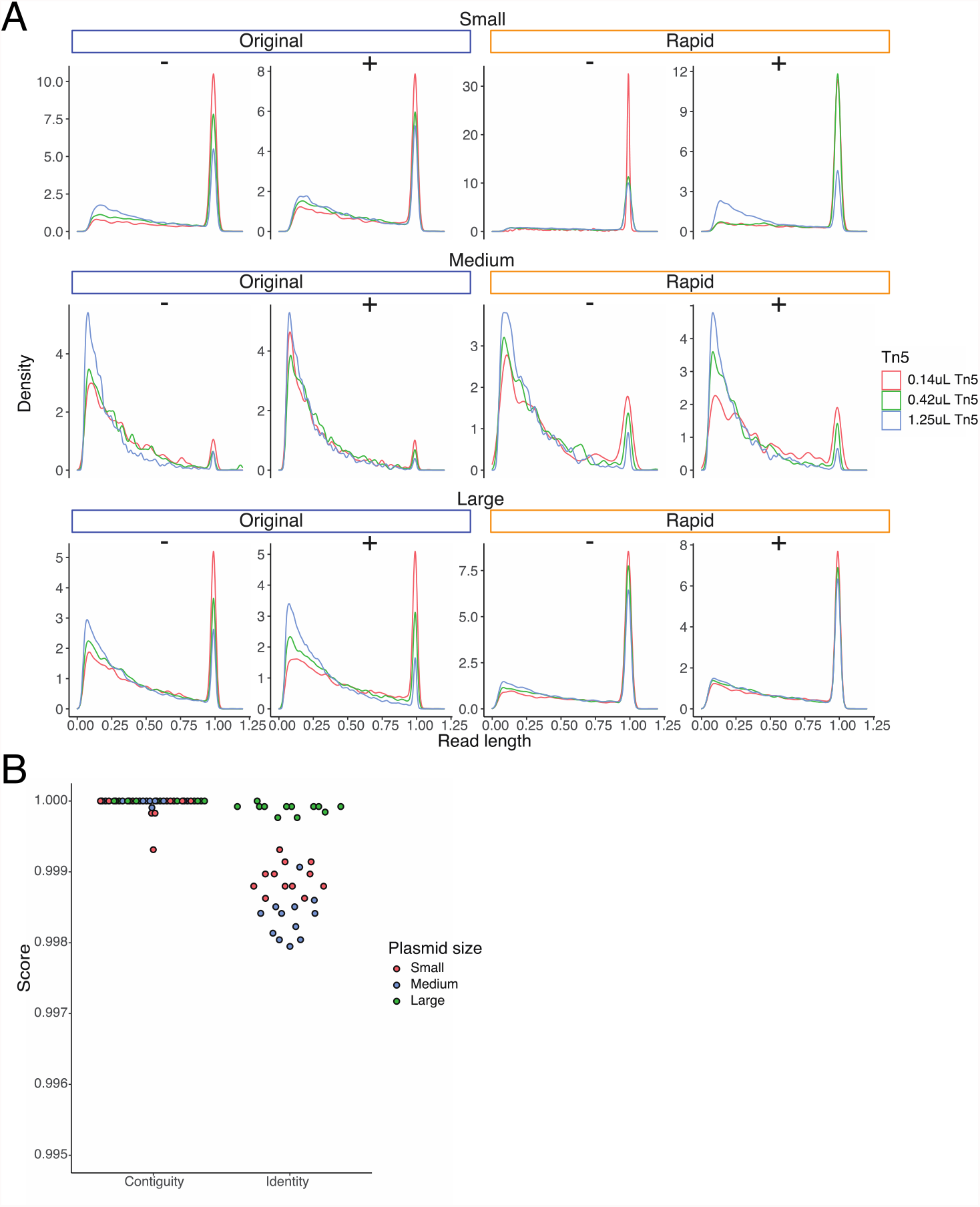
**(A)**. To determine conditions that would maximize read length we processed three plasmids of various lengths (6kbp to 13kbp) through a modified pipeline. Density plots of read lengths, normalized to the full length plasmid, are shown under the conditions of changing length (top to bottom), amount of Tn5 (colors), the duration of tagmentation (left and right), as well as method for inactivation (-for SDS only, + for SDS with heat inactivation). **(B)**. Once the reads from each of the conditions were assembled we calculated contiguity and identity scores for each assembly. Identity scores cluster by plasmid type due to sequence complexity and are unrelated to reaction conditions.

**Sup. figure 3.**
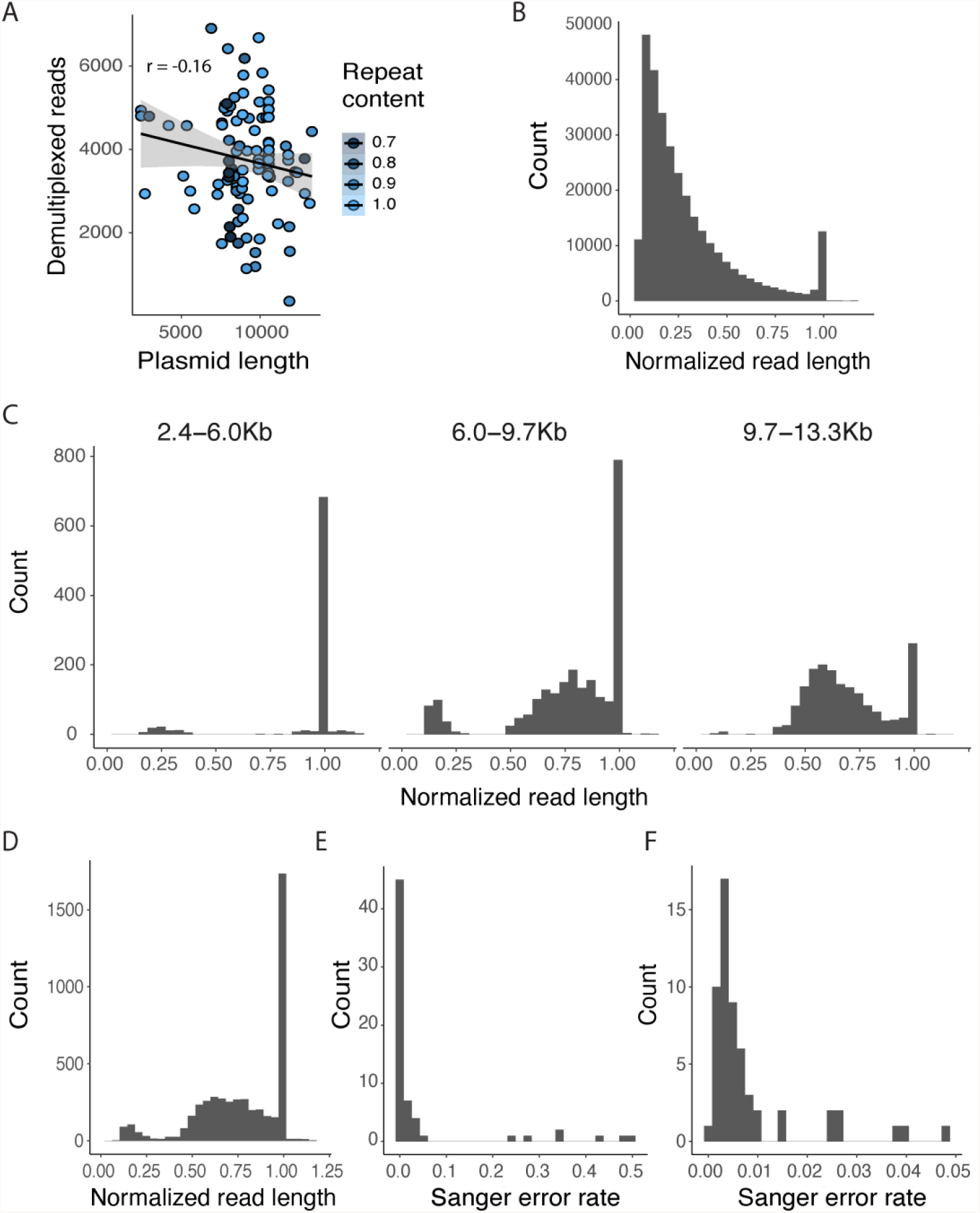
**(A)**. The correlation of recovered Nanopore sequencing read counts to input plasmid length, where point color indicates the relative repeat content of the input plasmid. **(B)** Histogram of unfiltered Nanopore read lengths normalized to the originating plasmid sequence length. **(C)** Post-filtering, normalized read lengths across plasmid lengths and **(D)** the global distribution. **(E)** Error rates for all 64 Sanger sequencing trace files after quality control and **(F)** after removing 7 traces with error rates over 10%.

**Supplemental figure 4.**
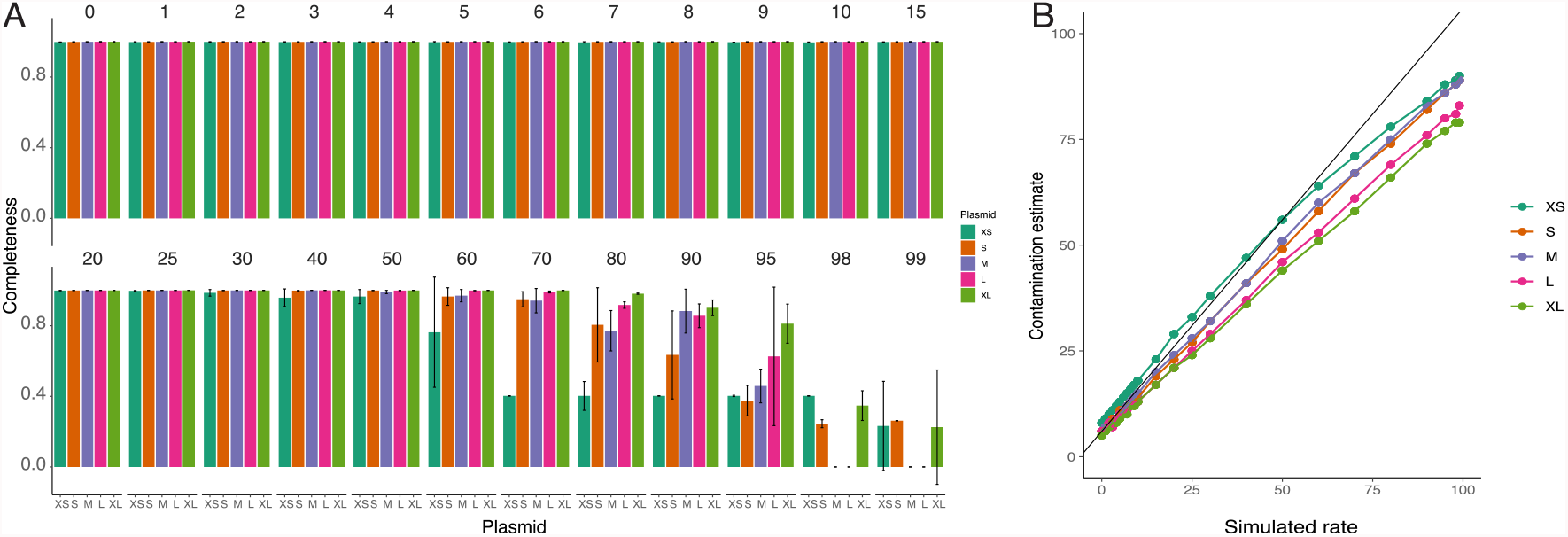
**(A)** Plasmid assembly completeness from simulated reads. Each of five plasmids (XS=extra small, S=small, M=medium, L=large, XL=extra large) were mixed with randomly sampled reads from the remaining 91 plasmids at contamination levels from 0% to 99%. Assembly issues in the smaller plasmids started at 30-40% contaminated, with most assemblies having serious issues at contamination levels greater than 80%. **(B)** Contamination estimates when reads are mapped against the known plasmid map instead of the assembled reference as in figure 3. Here contamination estimates better track simulated rates to the upper limits of contamination.

